# Architecture and rearrangements of a sperm-specific Na^+^/H^+^ exchanger

**DOI:** 10.1101/2023.10.04.560940

**Authors:** Kamalendu Pal, Sandipan Chowdhury

## Abstract

The sperm-specific sodium hydrogen exchanger, SLC9C1, underlies hyperpolarization and cyclic nucleotide stimulated proton fluxes across sperm membranes and regulates their hyperactivated motility. SLC9C1 is the first known instance of an ion transporter that uses a canonical voltage-sensing domain (VSD) and an evolutionarily conserved cyclic nucleotide binding domain (CNBD) to influence the dynamics of its ion-exchange domain (ED). The structural organization of this ‘tripartite transporter’ and the mechanisms whereby it integrates physical (membrane voltage) and chemical (cyclic nucleotide) cues are unknown. In this study, we use single particle cryo-electron microscopy to determine structures of a metazoan SLC9C1 in different conformational states. We find that the three structural domains are uniquely organized around a distinct ring-shaped scaffold that we call the ‘allosteric ring domain’ or ARD. The ARD undergoes coupled proton-dependent rearrangements with the ED and acts as a ‘signaling hub’ enabling allosteric communication between the key functional modules of sp9C1. We demonstrate that binding of cAMP causes large conformational changes in the cytoplasmic domains and disrupts key ARD-linked interfaces. We propose that these structural changes rescue the transmembrane domains from an auto-inhibited state and facilitate their functional dynamics. Our study provides a structural framework to understand and further probe electrochemical linkage in SLC9C1.

## Introduction

Hyperactivation of sperm in female reproductive tracts is associated with dramatic changes in the flagellar dynamics, which is pivotal for fertilization^1^. The process involves key signaling events at the sperm membranes, such as calcium influx and membrane hyperpolarization, and in the cytoplasm, such as elevation of cAMP levels and pH. Several sperm-specific biomolecular machineries are critical for temporal and spatial coordination of these signal cascade events. For instance, the CatSper channels^2,3^ mediate calcium entry into the cell that trigger phosphorylation events accompanying sperm motility changes. Another unique protein which has coevolved with CatSper in metazoans and is a critical player in sperm capacitation is the protein product of the SLC9C gene^4–6^.

The SLC9 family of ion exchangers are membrane proteins that couple the uptake of Na^+^ ions to proton extrusion and play key roles in salt and pH homeostasis in all kingdoms of life^7,8^. Mammals express thirteen different subtypes of SLC9s which are organized into three evolutionarily distinct sub-families – 9A, 9B and 9C^9^. Some of them, such as SLC9A1, are ubiquitously expressed at the plasma membrane, where they fulfill critical house-keeping functions. Others, such as SLC9A9 exhibit tissue and organelle-specific expression and accomplish specialized functions centering around their abilities to maintain salt and pH balance in specific cellular compartments. The role of the sperm-specific SLC9C1 in sperm physiology was first established using knock-out mouse models^10^ which showed that SLC9C1-null mice were infertile and showed severely compromised sperm motility. More recently a genetic mutation in human SLC9C1 has been linked to asthenozoospermia^11^. SLC9C has been recognized to be a potential target for the treatment of male infertility as well as the design of non-hormonal male contraceptives^12^.

SLC9C exhibit characteristic differences from other SLC9s in terms of their structural and regulatory attributes. In SLC9As and 9Bs, and their prokaryotic orthologs of known structure, the membrane-delimited ion-exchange domains (EDs) are tethered to relatively short or largely unstructured C-terminal soluble domains^13–18^. In contrast, SLC9Cs, feature a homologous ED, a canonical voltage-sensing domain (VSD) and a cyclic nucleotide binding domain (CNBD), interconnected via long, structured linkers, all within a single polypeptide. Foundational experiments recently performed with the sea urchin ortholog of SLC9C1^19^ (sp9C1) have revealed that it is a *bona-fide* electroneutral sodium-hydrogen exchanger. However, its ion exchange activity is under tight control of the VSD and is turned on only upon membrane hyperpolarization. cAMP binding tunes the voltage range over which it is functionally active, shifting it to less hyperpolarizing voltages. In terms of such regulatory mechanisms, sp9C1 is reminiscent of the well-studied HCN channels^20^. However, the mechanisms underlying voltage and cyclic nucleotide modulation of SLC9C1 function are not understood. To this end, we elucidate several structures of sp9C1 in different conformational states, using single particle cryo-electron microscropy (cryoEM) and reconstruction methods.

### Architecture and assembly of SLC9C1

We expressed full-length sp9C1 in mammalian cells as an eGFP fusion construct. Purification of the protein in digitonin micelles yielded biochemically monodisperse protein which was used for single particle cryoEM imaging (**Extended Fig. 1a-c**). We will first describe the overall architecture of sp9C1 obtained at pH 8 with K^+^ as the dominant cation which yielded a final reconstruction map at GS-FSC resolution of 3.8 Å (**Extended Fig. 1d, e**)

**Figure 1.**
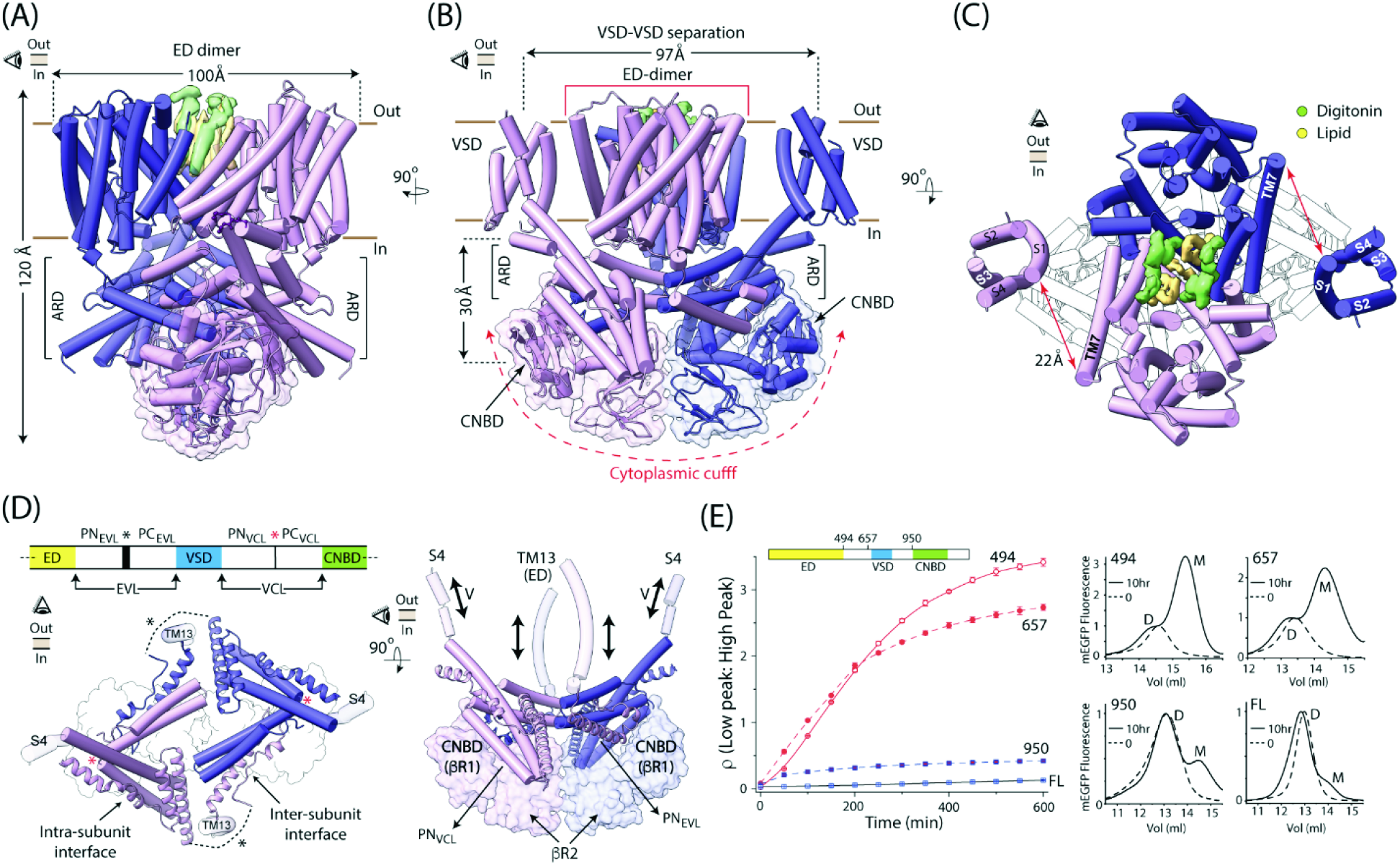
Architecture of *Strongylocentrotus purpuratus* SLC9C1. **A, B** Orthogonal views of the dimeric model of sp9C1, viewed parallel to membrane. The two subunits are colored in light purple and blue. Solid brown lines indicate the outer and inner membrane leaflets. VSD helices are hidden in **A** for clarity. **C** Top view of the sp9C1 along the membrane normal showing the intra-subunit separation between the VSD and ED and inter-subunit crevice of the EDs filled with lipid (yellow) and detergent (green) molecules. **D** *Left*, Top view of the ARD and parts of the protein which contribute to it. Protein segments from the same subunit are colored similarly (either light purple or blue). The Exchanger-Voltage-sensor Linker (EVL) and the Voltage-sensor-CNBD Linker (VCL) of both subunits are shown in cartoon and tube respectively. Black and Red * marks the flexible junction separating the two paddles of EVL and VCL respectively. Dotted line represents a region missing in density (corresponding to amino acid residues 535-570). Surface-outline of the cytoplasmic cuff, below the ARD, is also shown. *Right*, Side view showing the configuration of the ARD relative to the cytoplasmic cuff (shown as surface). PN_EVL_ of one subunit interacts with the CNBD of the other while PN_VCL_ interacts with the CNBD of its own subunit. ARD directly connects to the dynamic helices in VSD and ED (S4 and TM13 respectively) **E.** *Left*, Comparison of the time-dependent change of the ratio of low:high molecular weight species fraction (putatively monomer:dimer) under mildly destabilizing conditions for full-length sp9C1 (FL) and different truncation mutants. *Right,* Representative FSEC profiles of the proteins at the beginning and end (10hr) of the disassembly period, with M and D marking the monomer and dimer species for each.

The EDs of the two subunits pack against each other (**Figs. 1a-c, Supplementary Video 1**) forming the core of the protein. Each ED comprises 13 transmembrane helices (TM1 through TM13), organized in a manner similar to other members of the SLC9 family (**Extended Fig. 2a, b**). Seven of the thirteen transmembrane helices (TMs 1-3 and 7-10) form the ‘dimerization domain’ and the remaining six (TMs 4-6 and 11-13) constitute the ‘core domain’ (**Extended Fig. 2c**). A wide, cytoplasmically-accessible funnel, lined with negatively charged residues, is observed between the dimerization and core domains of the ED (**Extended Fig. 2c and d**). The tip of the funnel features conserved residues poised to bind a Na^+^ ion^21^. Thus, in our structure the ED is in an “inward open” state. Despite the structural similarity at an individual subunit level, the dimeric organization of the two EDs in sp9C1 is markedly different from other SLC9s (**Extended Fig. 2e**). TM10 of the two subunits of sp9C1, which line the inter-subunit cavity, tilt further away from each other by ∼12° causing their extracellular ends to splay apart by an additional ∼12Å, thus widening the crevice at the ED-dimer interface. In all our reconstructions, we were able to reliably identify densities for three lipid and four detergent molecules in this extracellularly accessible cavity (**Fig. 1c**). Lipid occupancy of this cavity might determine the stability of the sp9C1 dimers and act as clamps, holding the dimerization domain in place as the core helices move during the gating cycle of the ED^15,18,22,23^.

**Figure 2.**
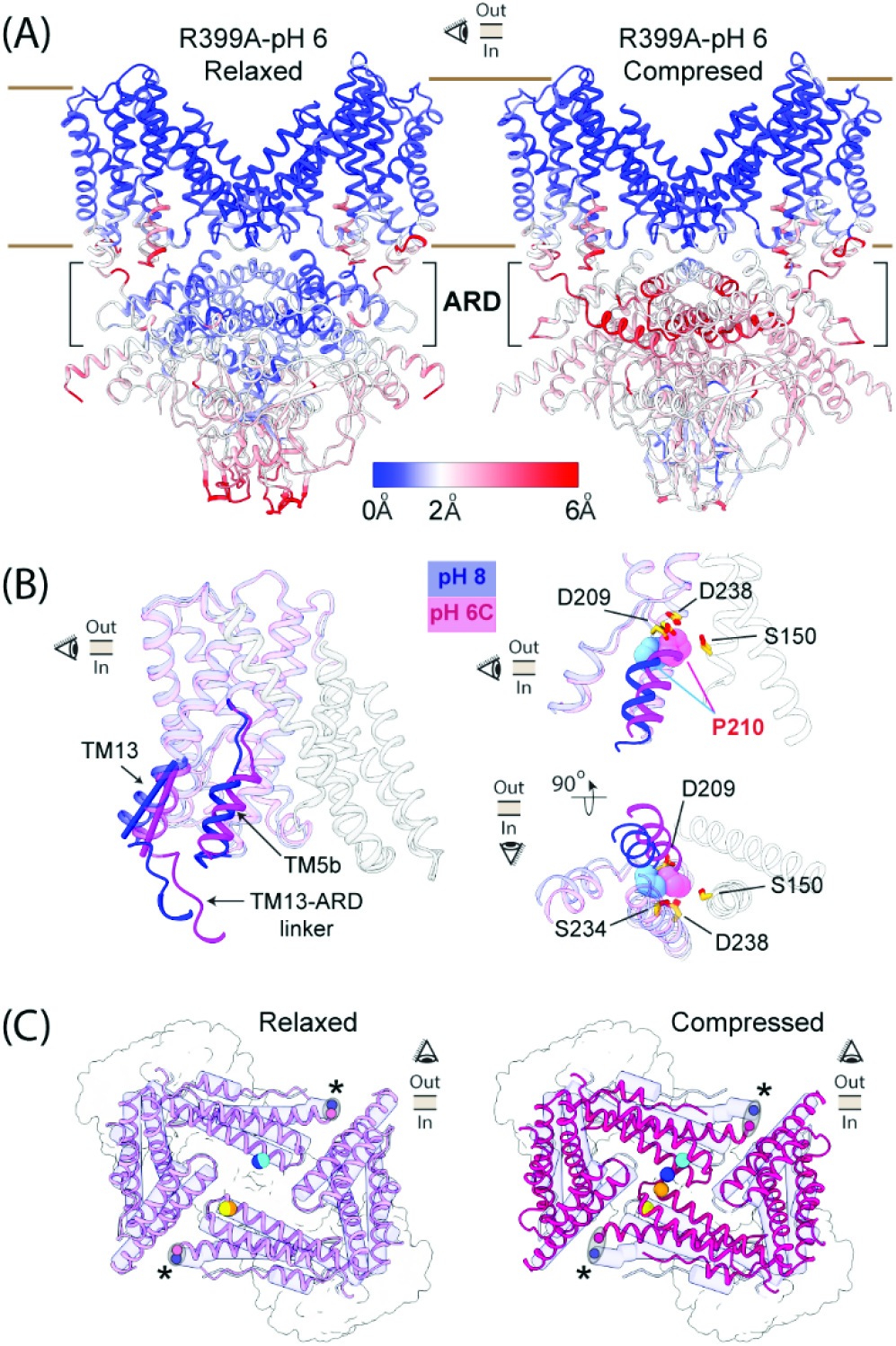
Coupled pH dependent rearrangements between ED and ARD in the R399A mutant of sp9C1. **A** Side views of the two structural classes of sp9C1-R399A at pH 6 (*Left*, Relaxed class and *Right*, Compressed class), looking along the short axis of the ED. Different parts of the protein are colored according to displacements of the Cα residues, with respect to the R399A-pH 8 structure. VSDs were omitted in the final models for the compressed or relaxed classes. **B** *Left*, Side view of a single subunit of the ED of R399A at pH 8 (blue) and pH 6-Compressed class (light magenta) showing the displacements of TM5b, TM13 and the short linker connecting TM13 and ARD. Solid cylinders highlight the helical axes of the intracellular ends of TM13. *Right*, Close-up of the Na+ binding site in the two conformations, in a side-view (top) or a bottom-up view (bottom). Residues forming the ion-binding site (in sticks) remain unchanged between pH 8 and pH 6, while P210 (spheres) is pushed into binding site due to the movement of TM5b. **C** Top view of the ARD in the R399A-pH6 Relaxed Class (*Left*) and R399A-pH6 Compressed class (*Right*) laid above the cytoplasmic cuff (surface outline). ARDs of the pH 6 are shown in pink and magenta while that of the pH 8 is shown in blue transparent tubes. Cyan and blue spheres indicate the Cβ atoms of H914 in a single subunit of R399A-pH8 and R399A-pH 6 structures respectively. Similarly, yellow and orange spheres mark the Cβ atoms of Q917 in pH 8 and 6 respectively. Relative to pH 8, there is significant rearrangement in the compressed class (*right*) but not in the relaxed class (*left*). The two ellipses and their foci (marked by *) indicate the displacements of Cα of residue 488, in the TM13-ARD linker.

The VSDs (each comprising helices S1 through S4) are structurally estranged from the EDs, appearing as “floating buoys” on either side of the ED-dimer interface. At the closest, the VSD (extracellular end of S1) is ∼22Å away from the ED (extracellular end of TM7) of the same subunit (**Fig. 1c**). This arrangement is in striking contrast to VGICs where the VSDs form intimate contacts with the ion translocating pore domain and the shared interfaces profoundly impact voltage-dependent channel opening^24^. The lack of any direct contact between the VSD and the ED in sp9C1 indicates that the allosteric mechanism underlying its voltage-regulation is different from that in VGICs. It is noteworthy that while the local resolution of our density map (FSC = 0.5) reaches ∼3.2Å in the core of the protein, it drops to 4-6Å in the peripheral regions, including the VSD (**Extended Fig. 1e**). This is probably due to its lose packing with the protein core which leads to its higher structural flexibility. Nevertheless we were able to reliably trace the backbone of most of the VSD helices (**Extended Fig. 3a, b**). The S4 helix of sp9C1-VSD, which is critical for its voltage-sensitivity, has six regularly spaced positively charged residues. The intracellular end of S4 forms a 9-residue-long 3_10_-helix harboring three positive charges (R809, R812, K815) which are highly conserved in different sp9C1 orthologs (**Extended Fig. 3c**). The remaining three positive charges (K800, R803 and R806) are arranged on the extracellular, α-helical end of S4. These three residues are relatively less conserved, particularly in the mammalian variants of SLC9C1, which could contribute to functional divergence in their voltage-sensitivities. The limited resolution of the VSD precludes unambiguous deduction of its conformational state. However, given the strong hyperpolarizing voltages necessary to drive the sp9C1-VSD into its resting (or down) state^19^, it is likely that in our structure (at 0mV) it is in the activated (or up) conformation.

**Figure 3.**
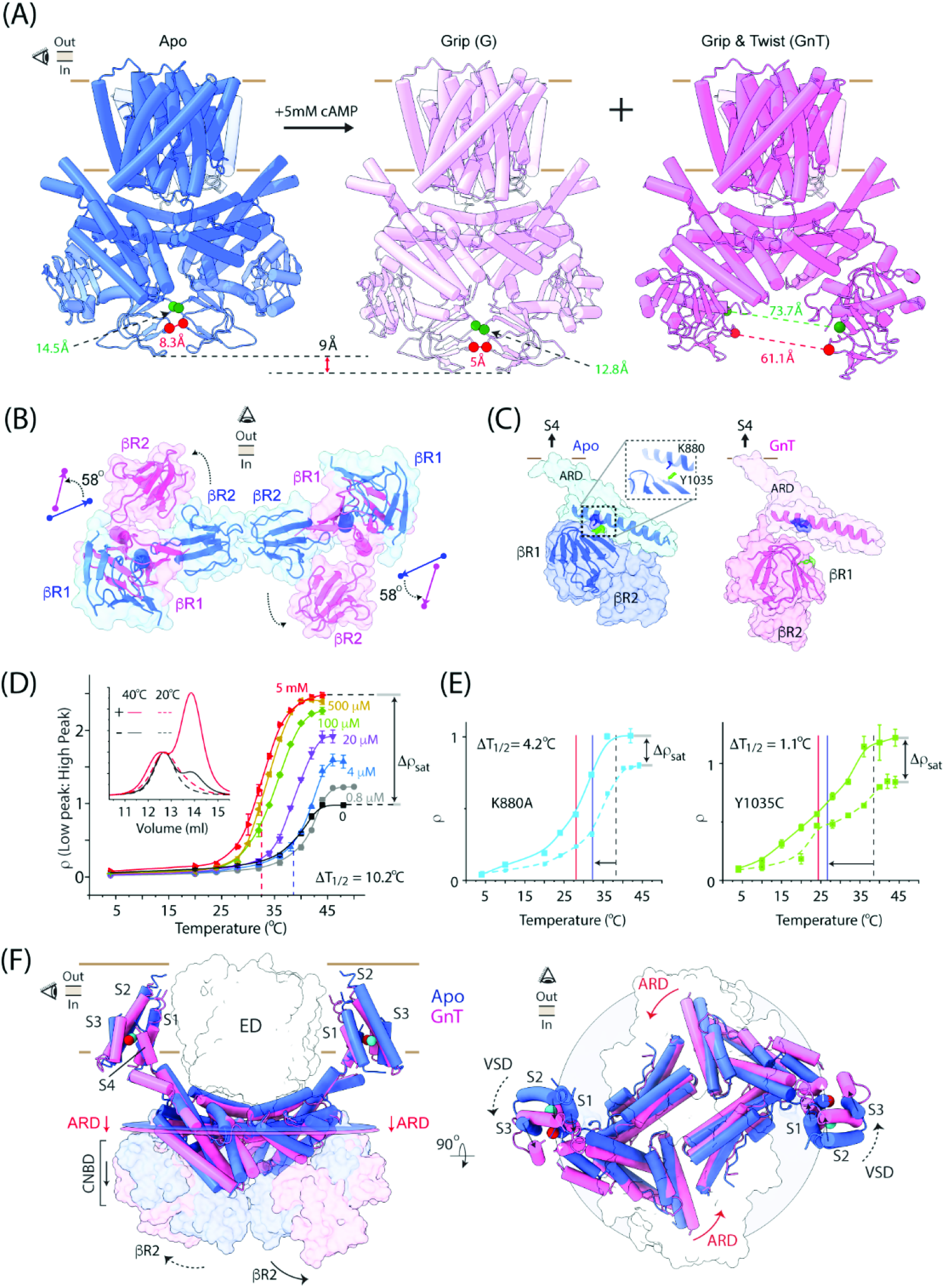
cAMP dependent rearrangements in sp9C1. **A** Side views of cAMP-free (Apo) sp9C1 (pH 6/NaCl model) and two cAMP bound classes, Grip (G) in pink and Grip-and-Twist (GnT) in dark pink, looking along the long axis of the ED. Between the Apo and Grip, the cytoplasmic cuff remains closed but moves downward by ∼9Å. Red and Green spheres indicate residues Thr1170 and Ala1140 in the βR2 domain of the two subunits which separate dramatically in the GnT class. **B** Top-down views of the cytoplasmic cuff in the Apo (blue, pH 6/NaCl structure) and GnT (dark pink) states showing the ∼58° twist of the axis connecting the center of masses of βR1 and βR2. **C** Side view of PN_VCL_ segment of the ARD and the cytoplasmic domains of a single subunit showing the interface between them in the Apo state (blue, pH 6/NaCl structure) which is disrupted upon cAMP binding in the GnT class (dark pink). Two conserved residues at this interface, K880 and Y1035, are highlighted in blue and green respectively. **D** TDCs of WT sp9C1 at different cAMP concentrations. Vertical dashed lines represent the T_1/2_ of sp9C1 in cAMP-free (black) and 5mM cAMP (red) conditions. Inset shows the chromatography files of sp9C1 incubated at 20°C (dotted) or 40°C (solid), in presence (red) or absence (black) of cAMP (indicated by + and – signs respectively). **E** TDCs of K880A and Y1035C mutants of sp9C1 in cAMP-free (dotted curve) or 5mM cAMP (solid curve) conditions. Vertical dotted line indicates the T_1/2_ of WT sp9C1 in absence of cAMP. Blue and red vertical lines indicate the T_1/2_ of the respective mutants in absence and presence of 5mM cAMP, respectively. **F** Overlay of the GnT class (dark pink) and Apo (blue, pH 6/KCl structure) showing the displacements of the ARD and the discernible parts of the VSD. The circular planes represent the average planes of the ARD, colored according to the respective structures. Spheres represent the center of masses of the modeled parts of the VSD that are common to both structures. The cytoplasmic cuffs are shown as transparent surfaces.

In each sp9C1 subunit, the CNBD hangs ∼30Å below the inner leaflet of the membrane. It has an evolutionarily conserved fold comprising an 8-stranded β-jelly roll (βR1) flanked on the N-terminus by the αA helix and on the C-terminus by αB and αC helices. Unexpectedly, the CNBD is C-terminally linked to a second 8-stranded β-jelly roll (βR2) via an α-helical linker (αD helix) (**Extended Fig. 3d**). Extensive hydrophobic contacts and a possible cation-π interaction, mediated by conserved residues (W1091 and R1185), are observed at the interface between βR2 and αC helix (**Extended Fig. 3d,e**). The βR2s of the two subunits are structurally apposed against each other, and together with the CNBDs, form a “cytoplasmic cuff”. CNBDs of cyclic nucleotide modulated ion channels do not feature a comparable βR2 domain^25–27^. While the specific functional role of βR2 is unclear, we will presently discuss a possible role of the βR2 in guiding cyclic nucleotide dependent conformational changes in sp9C1.

A distinct scaffolding domain, which we refer to as the ‘allosteric ring domain’ (ARD), directs the three dimensional arrangement of the ED, VSD and CNBD in the context of the sp9C1 dimers (**Fig.1d**). The ARD fills the gap between the cytoplasmic surface of the transmembrane domains and the CNBDs. It is formed by the association of the polypeptide segments connecting the ED and VSD (the Exchanger-Voltage-sensor Linker or EVL) and the VSD and CNBD (the Voltage-sensor-CNBD linker or VCL). Each of the linkers are shaped like two paddles, annotated as PN_<X>_ or PC_<X>_ (where N and C designate the N-terminal or C-terminal paddle and X is either EVL or VCL) connected by a flexible hinge. Inter-subunit interactions between PN_EVL_ and PC_VCL_ and intra-subunit contacts between PC_EVL_ and PN_VCL_ drive the assembly of the ARD. In addition, the ARD interacts intimately with the βR1 of CNBDs – PN_EVL_ is nestled in a groove formed between PC_VCL_ and βR1 of the neighboring subunit while PN_VCL_ rests atop of the βR1 of the same subunit (**Fig. 1d** **and Supplementary Video 1**). The overall arrangement of the cytoplasmic domains and ARD of sp9C1 is similar to the SOS1 transporter^17^, which is a plant ortholog of SLC9s, although the latter does not feature a VSD and is not regulated by cAMP.

The characteristic arrangement of the ARD suggests that it may play an important role in the assembly of the sp9C1 dimers. To test this, we compared the stabilities of full length (FL) sp9C1 with 3 truncation mutants (named 950, 657 and 494 referring to the amino acid positions of the truncation sites) using Fluorescence Size Exclusion Chromatography (FSEC)^28,29^. All 4 proteins, expressed as eGFP fusion constructs and affinity purified in digitonin micelles, were largely monodisperse with retention volumes consistent with a dimeric assembly (**Fig.1e**). However, when exchanged into mildly destabilizing conditions, a second peak, corresponding to a monomer, was observed. Constructs truncated at 657 and 494 were drastically more unstable than FL-sp9C1 as reflected by a much faster and greater extent of disassembly over a 10hr period. In both these constructs the ARD is partly or entirely deleted. Thus the intra-and inter-subunit interactions at the level of the ARD are critical for stability of the sp9C1 dimers. The 950 construct, in which the CNBD, βR2 and the C-terminal tail was deleted, also exhibited elevated breakdown relative to the full-length protein. Thus the closed cuff arrangement of the cytoplasmic domains also contributes to dimeric stability, either via direct inter-subunit interactions between the βR2 domains or by facilitating a stable arrangement of the ARD via the ARD-CNBD interfaces.

### Coupling of pH dependent dynamics of the ED and the ARD

The ARD structurally interweaves the three key dynamic elements of sp9C1 through covalent and non-covalent interactions. This raises a compelling hypothesis that the ARD energetically couples their intrinsic rearrangements and might itself exhibit non-trivial relaxations in response to conformational changes in the ED, VSD and CNBD. To explore this possibility we pursued single particle reconstructions of sp9C1 under acidic conditions (pH 6). We reasoned that since protons are co-substrates of the ED, by elevating its concentration, we could ostensibly drive the ED into a different conformation and visualize how the remainder of the protein structurally adapts to it.

Our final maps of sp9C1 at pH6, with K^+^ or Na^+^ as the primary cation, reached GS-FSC resolutions of 3.7 Å and 4.2 Å respectively (**Extended Fig. 4a, b**). However the density for the VSDs worsened relative to the pH 8 reconstructions. We speculate that protonation of titratable residues in the VSD results in their increased structural heterogeneity, probably similar to VSD-mediated proton dependent inhibition of many VGICs^30–32^. Beyond this, very modest changes were observed in the remainder of the protein, such as 2-3Å displacements in the βR2 (**Extended Fig. 4c-f**). While it is possible that in digitonin micelles the ED is conformationally arrested, it is noteworthy that reconstructions of SLC9A1/CHP1 in lipid nanodiscs also show limited pH dependent rearrangements in the ED^16^, raising the possibility that the inward open state of the ED might be intrinsically favored in different SLC9s, particular under *in vitro* conditions. An alternate possibility is that the pKa-s of the titratable residues in the ED, which underlie its proton dependent regulation, is significantly lower than pH 6.

**Figure 4.**
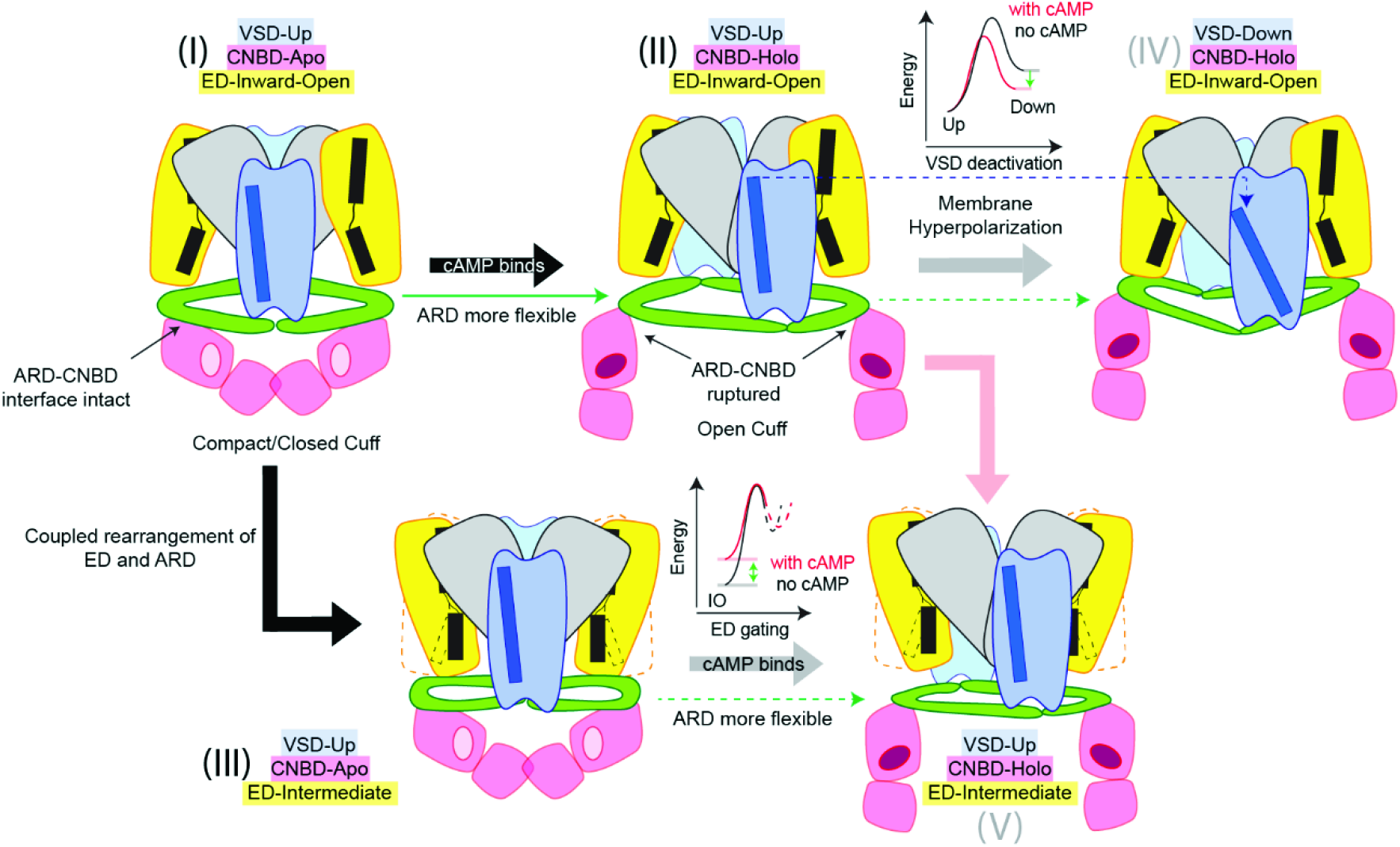
Proposed model for cAMP regulation of sp9C1. Abstract cartoons of sp9C1 in five distinct functional states are shown. In each case, core domains of ED are in yellow, the dimerization domains of ED are in gray, VSDs are in blue, ARD is in green and the cytoplasmic cuff is in red. TM5 of ED and S4 of VSD are illustrated in black and blue respectively. In state (I) (as represented by the structures of in pH 8/cAMP-free conditions), ARD is rigid and sp9C1 is inhibited by the compact form of the cytoplasmic cuff. Upon cAMP binding (II), the cuff opens and increases the flexibility of ARD. As the structural restraint imposed by the compact cytoplasmic is lifted, the ARD can undergo coupled rearrangements with VSD deactivation (IV) more easily and thus cAMP binding allosterically stabilizes the down state of the VSD. The ARD undergoes coupled rearrangements with ED (State (I) to (III)). This is also hindered by the compact cytoplasmic cuff through the ARD-linked interface. Upon cAMP binding, structural coupling between the ED and ARD becomes more favorable (State (III) relative to State (V)) and the energy barrier for the ED to transition out of the Inward-Open state decreases resulting in faster ion-turn over rates. Note that the structures of States (IV) and (V) have not been explored in this study.

Constant pH MD simulations of a prokaryotic electrogenic ortholog of SLC9s have suggested that a highly conserved positively charged residue (equivalent to R399 in sp9C1) plays a critical role in ion exchange, at least in part, by influencing the pKa of neighboring Asp/Glu residues which constitute the principle Na^+^ binding site (**Extended Fig. 5c**)^33,34^. Applying the same logic to sp9C1, we reasoned that neutralization of R399 might increase the pKa of nearby acidic residues (E233, D209, D238) and thereby facilitate capturing proton dependent conformational changes in the protein. Accordingly, we imaged the R399A mutant of sp9C1 at pH 8 and 6, with Na^+^ as the dominant cation. At pH8, reconstruction of the R399A mutant identified a single structural class with an overall GS-FSC resolution of ∼3.1Å (**Extended Fig. 5a**). However at pH 6 our reconstruction analyses revealed two distinct structural classes at GS-FSC resolutions of 3.5 Å and 3.1 Å which we refer to as the Relaxed (R) and Compressed (C) forms respectively (**Extended Fig. 5b**). While the R399A reconstructions recapitulated all key architectural features observed with WT sp9C1 (**Extended Fig. 5d**), they exhibit clear pH dependent rearrangements (**Fig. 2a, Supplementary Video 2**). Alignment of the pH8 and pH6 structures of the R399A revealed that upon acidification, in both the R and C classes, the TM5b moved upward and laterally by ∼2Å each (**Fig. 2b**). This movement causes the side-chain of P210 to be inserted into the substrate binding site, possibly altering protein-substrate interaction, although the sodium coordinating residues remain largely static (**Fig. 2b**). The last 2-3 turns of TM13 also undergoes a ∼12° change in tilt (**Fig. 2b**). The resultant RMSDs of these two regions of the protein between pH 8 and 6, is considerably larger with the R399A mutant than with WT sp9C1 (with K^+^ or Na^+^ as the dominant cation) (**Extended Fig. 5e**). At the level of the ARD however, dramatic changes were observed in Class C but not in Class R (**Fig. 2c**). In Class C, there is a clear displacement of the 6-residue long linker between TM13 and PN_EVL_ which culminates in an ∼8° inward rotation of the PN_EVL_. PC_VCL_ of the opposite subunit, tightly packed against the PN_EVL_, moves along with it as a rigid body (**Extended Fig. 5f**). As a result of this rotation, the PC_VCL_ of the two subunits which are 15.9Å apart (at level of residues H914 and Q917) in pH 8, are only 4.3Å apart in the pH 6 Compressed Class. These structural observations support our hypothesis that the ARD undergoes coupled rearrangements together with the ED. The ARD-βR1 interfaces however remain largely preserved between the pH8 and the pH6-compressed form (**Extended Fig. 5g**), indicating that the compact cytoplasmic-cuff moves with the ARD pointing towards the stability of the ARD-βR1 interfaces.

Functional measurements of sp9C1 indicate that the R399A mutation causes a dramatic decrease in ion turnover rates^19^. R399 is engaged with E233 via a salt-bridge interaction. This interaction likely couples the core helices of the ED as it toggles between the inward-open and the outward open states as observed in the NHE1-CHP1 complex^16^. The loss of function in R399A might arise from stabilization of an intermediate state of the transporter. Without the salt-bridge, the core helices get trapped in this state and are unable to isomerize into the outward open state. In addition, the R399A mutation in sp9C1 has also been shown to perturb VSD dynamics, causing the gating-charge displacement *vs* voltage curve to shift rightward along the voltage-axis and become shallower^19^. The lack of clear density for the VSD at pH 6 (much like what we observed with WT sp9C1) does not allow us structurally define such an intermediate state of the VSD. However, the coupled rearrangement of the ARD with the ED enables us to propose that the allosteric communication between the VSD and ED is mediated, at least in part, by the ARD.

### cAMP dependent re-arrangements in sp9C1

cAMP exerts two main effects on sp9C1 function. It reduces the extent of hyperpolarization required to activate the ion-exchange activity and stabilizes the resting or down state of the VSD^19^. We investigated the structural mechanisms underlying cAMP dependent modulation of sp9C1 via single-particle reconstructions of sp9C1 in presence of cAMP. Initial reconstruction analyses indicated that, in presence of cAMP, sp9C1 was structurally heterogenous and the heterogeneity was worse at pH 8 than at pH 6. Thus we pursued our reconstruction efforts of sp9C1, in presence of cAMP, only at pH 6. We were able to identify three distinct, almost equally populated, structural classes in the presence of cAMP (**Extended Fig. 6a**). The classes, which we call G (Grip), GnT (Grip and Twist) and GnTL (Grip and Twist Like) (**Fig. 3a** and **Extended Fig. 6b**) were resolved at GS-FSC resolutions of 4.1Å, 3.9Å and 4.2Å respectively. Much like our pH 6 reconstructions in cAMP free conditions, densities for the VSDs were not clear in the G and GnTL classes. In the GnT class, several intracellular helical turns of the four VSD helices could be resolved, which defined the position of the VSD within the plane of the membrane. The EDs of all three structural classes was practically identical to that observed in cAMP free state (**Extended Fig. 6c**). The cytoplasmic domains however exhibited large rearrangements.

When compared to the cAMP free structures, we find that in class G, the cytoplasmic cuff retains its compact/closed form but moves downward by about 9Å (**Fig. 3a**). In the GnT and GnTL classes however, the CNBD and βR2 domains of each subunit twist outward by ∼58°, opening the cytoplasmic cuff (**Fig. 3a, b**). As a result of this movement, the separation between the βR2 domains in the GnT and GnTL conformations increases dramatically by about ∼ 60Å, relative to the Apo and G conformations. For the GnT class, the density map was of sufficient quality to identify a cAMP molecule, bound in an anti-configuration (**Extended Fig. 6d**), within the CNBD. cAMP binding causes the bR1 and aC helix of the CNBD to move in closer to each other by ∼5Å. A similar conformational change in also observed in the CNBDs of classes G and GnTL (**Extended Fig. 6c**), although the cAMP density is relatively less clear. Thus, G, GnT and GnTL all represent distinct cAMP bound quaternary conformations of sp9C1.

Structural alignment of the cytosolic domains (CNBD and βR2) of the cAMP bound conformations with that of the apo form shows that the interface between the αC and βR2 remains relatively invariant during ligand binding (**Extended Fig. 6e**). Thus it is more likely that cAMP is gripped by the CNBD as a result of the movement of βR1 towards αC, as opposed to a movement of the αC-βR2 unit towards βR1. The movement of αC with respect to βR2 is possibly restricted by the hydrophobic interface between them. Thus βR2 might be acting as a structural rudder guiding the specific rearrangement of the CNBD. As βR1 moves inward to grip cAMP, the intra-subunit interface between βR1 and ARD interface is ruptured. This causes the cytoplasmic domain of each subunit to become relatively untethered to the rest of the protein and is free to move, which leads to the large conformational changes observed the GnT and GnTL states.

cAMP binding induced structural changes in the cytoplasmic cuff and rearranges the ARD. In the GnT conformation, the average plane of the ARD is displaced downward by ∼2Å with respect to the Apo state (**Fig. 3f**). Interestingly the PC_EVL_ and PN_VCL_ paddles rotate counter-clockwise about the membrane normal by ∼7°. The latter movement is particularly significant, since it leads to a robust, 9Å lateral displacement of the VSD helices within the plane of the membrane (**Fig. 3f, Supplementary Video 3**).

Since the closed cuff stabilizes the sp9C1 dimers, we gauged the effect of cAMP binding on the biochemical stability of sp9C1 dimers using a thermal disassembly assay. Purified sp9C1-eGFP was heated to different temperatures in absence or presence of cAMP (at different concentrations) and the monodispersity of the sample was analyzed using FSEC. A distinct lower molecular weight peak, likely corresponding to a disassembled monomer, became progressively more dominant with increasing temperature, saturating by ∼46°C. The ratio of the two species of distinct sizes (ρ) was plotted against the temperature to obtain a temperature-dependent disassembly curve (TDC) at each cAMP concentration. With increasing concentrations of cAMP, the saturating level of ρ (ρ_sat_) increased and T_1/2_ of the TDC decreased (**Fig. 3d**). Between cAMP-free and 5mM cAMP, ρ_sat_ increases from ∼0.9 to 2.4 and T_1/2_ reduces from 38°C to 28°C, indicating a large loss of dimer stability upon cAMP binding. A dose-response relationship for cAMP, based on ρ_sat_ or T_1/2_, indicates that cAMP interacts with purified sp9C1 with an EC_50_ of ∼12-15uM.

To determine the effect of CNBD-ARD interface on cAMP induced loss of protein stability, we mutated two conserved residues localized at this interface and determined their TDCs in the absence and presence (5mM) of cAMP. Y1035C and K880A destabilize the sp9C1 dimers in cAMP free-conditions, as reflected by a significantly lower T_1/2_ with respect to the WT. Both mutations also robustly decrease cAMP dependent effects (**Fig. 3e**) – Δρ_sat_ (the change in ρ_sat_ between cAMP-free and 5mM cAMP conditions) for the R880A and Y1035C mutants are ∼70% lower than that for the WT sp9C1 and there is a pronounced decrease in the cAMP dependent shift (ΔT_1/2_) of the TDCs. A likely interpretation of these observations is that Y1035C and K880A destabilizes the intra-subunit interface between the ARD (PN_VCL_) and βR1, thereby perturbing the compact closed-cuff arrangement of the cytoplasmic domains in the cAMP-free state. As a result, the cAMP dependent loss of stability caused by the opening of the cytoplasmic cuff is diminished in the mutants. Overall, the results of these thermostability experiments are consistent with our structural observations and points to an important role for the CNBD-ARD interface in governing the stability of the compact form of the cytoplasmic cuff, which in turn affects the cAMP dependent dynamics of sp9C1.

### Mechanism of electrochemical linkage in SLC9C1 and conclusion

Our reconstructions of sp9C1 in multiple conformations enables us to propose a biophysical basis underlying cAMP regulation of sp9C1 function (**Fig. 4**). We hypothesize that in the absence of cAMP and at depolarizing voltages, sp9C1 is in an auto-inhibited state. The inhibition of the ED arises from the specific arrangement of the ARD, which is in turn spatially constrained by the activated VSD (due to the rigid link between the S4 and PN_VCL_) and the apo CNBD (via the intra-and inter-subunit interfaces). Upon binding cAMP, the cytoplasmic cuff opens and relieves the structural constraint imposed by the cytoplasmic domains on the ARD. This enables the ARD to rearrange more easily as the voltage-sensor deactivates (S4 moves down), thereby decreasing the hyperpolarizing voltages required to drive the voltage-sensor into its down state. Concurrently, it decreases the energy barrier for the ARD to undergo coupled rearrangements with the ED, as it toggles between its inward-open and outward-open states, catalyzing ion exchange. Overall, cAMP induced potentiation of sp9C1 ion turnover rates and facilitation of voltage-sensor deactivation proceeds via relieving the transporter from an auto-inhibited state.

While further studies will be necessary to etch the energetic and structural landscape underlying SLC9C1 function it is noteworthy that our model of cAMP dependent facilitation of SLC9C1 activity bears striking similarities to that proposed for HCN channels^35,36^. In the case of the latter, the VSDs are at best weakly coupled to cAMP binding^37^ and the scale of cAMP dependent rearrangements is small^27,38^ as compared to what we observe here with SLC9C1. Yet it is remarkable how, despite architectural and dynamic differences, the underlying biophysical themes underlying cAMP regulation are convergent between the HCN channel and SLC9C1, arguably an HCN transporter.

## Methods

### Expression and purification

The cDNA fragment encoding full-length *Strongylocentrotus purpuratus* SLC9C1 was synthesized (Genscript Inc.) and cloned into a modified pEG BacMam vector^39^, with a PreScission protease cleavage site followed by a C-terminal mEGFP-Twinstrep tag. All mutants of sp9C1 (R399A and the constructs truncated at amino acid positions 494, 657 and 950) were generated in the background of this parent construct using standard molecular biology techniques (Genscript Inc.). For protein expression, HEK293F cells, cultured in suspension in Freestyle 293 media (supplemented with 2% Heat Inactivated FBS), were transfected with plasmid DNA, isolated from large volumes of bacterial cultures using Endotoxin free Plasmid Purification kits (Qiagen). Linear PEI (25kDa) was used for transient transfections at a ratio of 1:3 (plasmid:PEI mass ratio). Post-transfection, cells were grown for 12-14 hrs at 37°C, following which sodium butyrate was added to the transfected cells to a final concentration of 10mM and the cultures were transferred to 30°C and grown for another 48-54 hrs. Cells were collected by centrifugation at 3,000g for 30mins and washed with PBS. Pelleted cells were flash frozen in liquid nitrogen and stored at −80°C until use.

For purification, frozen cell pellets were resuspended in chilled lysis buffer (300mM NaCl, 50mM Tris, 10mM DTT, 20% glycerol, 1% digitonin, pH 8), briefly sonicated on ice and gently agitated at 4°C for 1-1.5 hrs. The whole-cell extracts were spun at 100,000g for 1 hr and the supernatant was incubated anti-GFP-nanobody resin (generated by PCF, University of Iowa) for 8-10 hrs. Protein bound resin was washed 4 times in batch mode, each time with 5 resin volumes of wash buffer (300mM NaCl, 50mM Tris, 10mM DTT, 10% glycerol, 0.1% digitonin, pH 8). After the last wash, the resin was resuspended in 2x resin volume of wash buffer and incubated with Precision protease (ThermoScientific) for 12-14hrs. The protein released by protease cleavage was concentrated to ∼500ul using 100kDa centrifugation filters and the resultant protein was further purified by gel filtration chromatography. The gel filtration buffer used was modified according to desired condition for single-particle imaging. For pH 8 samples, the buffer was 300mM NaCl, 25mM HEPES, 1mM TCEP, 0.05% digitonin, buffered to pH 8 using NaOH and for pH 6 samples, the buffer was 300mM NaCl, 25mM MES, 1mM TCEP, 0.05% digitonin, buffered to pH 6 using NaOH. For conditions where K^+^ was used instead of Na^+^, NaCl was replaced with KCl and KOH was used to bring the final pH of the solution to the desired value. 1-1.25ml of the peak fractions of the protein was collected and concentrated to ∼2.5-3.5mg/ml for preparing cryoEM grids.

For all thermostability experiments, the constructs were similarly expressed in HEK293F cells. After extraction in digitonin buffer, the protein was purified using Streptactin affinity resin (IBA Life Sciences). Resin bound protein was washed as before and eluted using wash buffer, supplemented with 10mM Desthiobiotin. Eluted protein was further purified by gel filtration chromatography using the SEC buffer: 300mM NaCl, 50mM Tris, 0.05% digitonin, buffered to pH 8. Peak protein fractions were combined and concentrated to ∼1.5mg/ml, flash frozen in liquid nitrogen (in 5µl aliquots) and stored in −80°C until use. All steps of protein purification were performed at 4°C.

### Thermostability assessment using Fluorescence Size Exclusion Chromatography

For all thermostability tests, the destabilizing buffer (TSB) was: 300mM NaCl, 50mM HEPES, 50mM MES, 1mM TCEP, 2mM DDM, 0.4mM CHS, buffered to pH 6. For the time-dependent disassembly experiments, a frozen aliquot of protein was thawed and diluted 200-fold (by volume) into the TSB and immediately injected (in under 1min) into the FSEC instrument (Shimadzu). Periodic injections (every 50min) of the sample were performed using the SIL40C autosampler and over the entire period the samples (diluted into TSB) were held at 4°C. For experiments probing cAMP dependent change in thermostability, thawed protein was diluted 200-fold into TSB, supplemented with an appropriate concentration of cAMP, incubated for 1min and heated to the desired temperature for 10min (on a PCR thermocycler) and the sample was subjected to FSEC analysis within 5mins of the heat step. In all cases, Superose 6 10/300 Column was used for separation of species and the FSEC buffer was: 300mM NaCl, 20mM Tris, 100µM GDN, pH 8. The column and buffers were at room temperature for all experiments. The chromatography profile of the different species was monitored by RF20Axs detector with Ex./Em. wavelength settings of 488nm/510nm. Each chromatography profile (between 10 and 16ml) was fitted to a sum of 2 Gaussians: 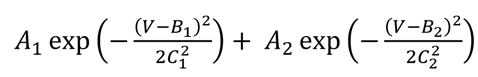. For all fits, B_1_ was constrained to be < B_2_ implying that the first gaussian term accounts for the higher molecular weight (dimer) species and the second term accounts for the disassembled monomer. The relative population of the 2 species was calculated as: A_2_C_2_/A_1_C_1_. In all cases, experiments were performed in duplicates with different batches of purified protein. All error bars are the standard deviation of measurements.

### CryoEM grid preparation and imaging

For all samples imaged under cAMP free conditions, 3.5ul of concentrated purified protein was applied to glow-discharged copper holey carbon grids. For the cAMP dataset, SEC purified protein was first concentrated to ∼1.5mg/ml, diluted 100-fold (volumetrically) into SEC buffer, supplemented with 2.5mM cAMP and incubated for 2hrs on ice. The protein was subsequently concentrated to ∼3mg/ml and an additional amount of cAMP was spiked into it so that the final protein and cAMP concentrations were ∼2.5mg/ml and 5mM respectively. The sample was incubated overnight on ice before preparing cryoEM grids. R 1.2/1.3 grids were used for all datasets, except for the pH 6/KCl dataset where R 2/2 grids were used. In all cases, grids were blotted for 5.5s at 100% humidity with a blot force of 0 and then plunge frozen in liquid ethane using a Vitrobot Mark IV (Thermofisher Scientific). All data were collected on a Titan Krios (electron microscope operating at an accelerating voltage of 300kV), equipped with a K3 Detector (Gatan). Images were recorded with EPU software (ThermoFisher Scientific) in super-resolution mode with a pixel size of 0.54 Å or 0.69 Å, and a nominal defocus of −0.9 to −1.9µm. For all datasets, a total dose of 65 electrons per Å^2^ was used, which was fractionated over 40 frames.

### Image Processing and Map Calculation

Image processing and map calculations were performed using CryoSPARC^40^ and Relion^41^. The recorded movies were motion corrected (Patch motion correction) and then subjected to contrast transfer function (CTF) estimation (Patch CTF). Micrographs with CTF resolution > 6Å or total pixel drift > 60 pixels were discarded. Blob picker was used to pick particles from 500-1000 images, which after several rounds of 2D class averaging, yielded reasonable 2D projections. These 2D classes were used for template based particle picking from curated movies. Furthermore, these (relatively few) particles were also subjected to Ab initio reconstruction (1 class) and subsequent homogenous refinement to generate a representative initial 3D map of the protein. The template picked particles were subject to several rounds of heterogenous refinements using the initial 3D map as reference. At each step of heterogenous refinement, the particles which get classified to low resolution classes were discarded. The set of particles thus retained were refined using Non-uniform^42^ and local refinement techniques on CryoSPARC (with C2 symmetry imposed), exported to Relion and further classified using 3D classification, without image alignment. The optimal set of particles were imported back into CryoSPARC and subject to CTF refinements^43^ to obtain the final maps. For processing of all datasets, except for R399A, particle stacks were binned by 2.

### Model Building and Refinement

An initial model of sp9C1 was generated using AlphaFold^44^ using its Colab platform. This initial model was divided into segments which were fitted into density maps and subsequently modified on COOT^45^. To generate the final models, several iterations of manual model building and real space refinement against full maps were performed in Phenix^46^. NAMDINATOR^47^ was used improve model building, particularly in instances where there are large domain level rearrangements. In 3 of the 10 cases, (namely 8K, 8Na and GnT) where the VSD densities are featured, the initial model building and refinement for the VSDs were done with the unsharpened map (because sharpening causes fragmentation of the densities in these regions) while the remainder of the model was generated using the sharpened maps. In all instances, the final map-to-model validation were performed against the sharpened maps. The final refined atomic models were validated using MolProbity^48^. All structural analyses were performed on UCSF Chimera^49^ or MATLAB (Mathworks). Structural figures were generated using UCSF ChimeraX or Pymol.

## DATA AVAILABILITY

All experimental data are available upon reasonable request. Sharpened, unsharpened and half-maps for each of the 10 reconstructions along with their PDB models have been deposited to EMDB/PDB repositories.

## ACKNOWLEDGEMENTS

This research was supported by grants to S.C. from NIH (R01-GM145719) and Department of Molecular Physiology and Biophysics, University of Iowa. We thank Dr. Vera Moiseenkova-Bell and acknowledge the use of instruments at the Beckman Center for Cryo-Electron Microscopy at the University of Pennsylvania Perelman School of Medicine. We thank Dr. Stefan Steimle for assistance with Krios microscope operation and cryoEM grid preparation at the Beckman Center for Cryo-Electron Microscopy at the University of Pennsylvania Perelman School of Medicine. We thank Drs. Nicholas Schnicker and Zhen Xu (PCF, University of Iowa) for providing anti-GFP conjugated resin for affinity purification of protein and Sankar Baruah (PCF, University of Iowa) for help with gel filtration chromatography.

## CONTRIBUTIONS

S.C. conceived and designed the study; S.C. and K.P. performed biochemistry experiments; S.C. performed single particle reconstructions; K.P. built atomic models; S.C. and K.P. prepared figures for manuscript; S.C. wrote the manuscript with inputs from K.P.

## COMPETING INTERESTS

The authors declare no competing financial interests.

---

**Extended Figure 1.**
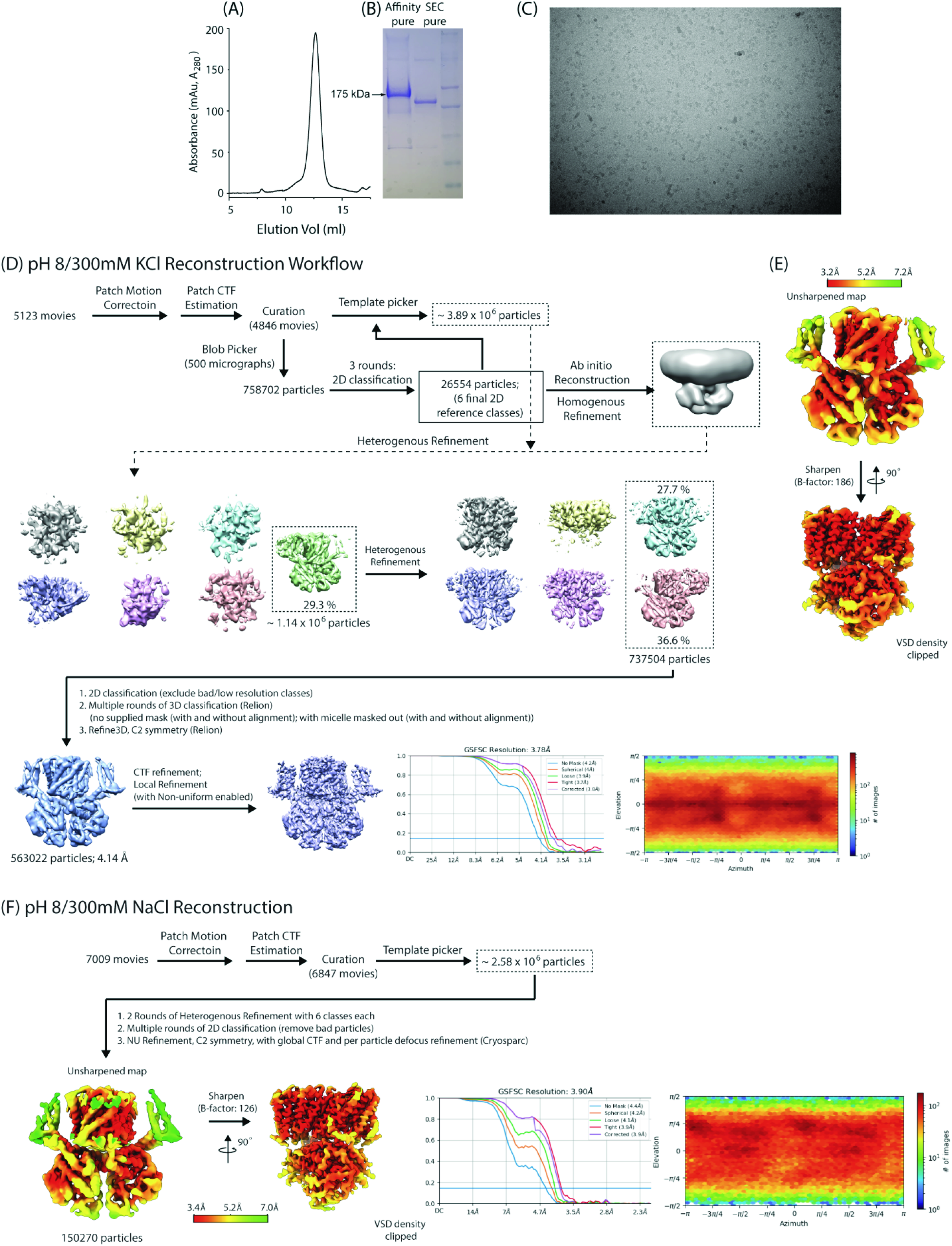
Purification of WT sp9C1 and reconstruction at pH 8. **A** Representative size exclusion chromatography profile of sp9C1 affinity purified using anti-GFP nanobody resin. The SEC Buffer was 300mM NaCl, 25mM HEPES, 1mM TCEP, 0.05% Digitonin, buffered to pH 8. **B** Representative SDS-PAGE gel of SEC purified sp9C1 obtained from streptactin affinity purification with mEGFP tag still intact (Lane A) and from anti-GFP nanobody affinity purification with a cleaved mEGFP tag (Lane B). (Lane C is the molecular weight marker). **C** An example micrograph of sp9C1 in pH 8 showing monodisperse single particles. **D** Single particle reconstruction workflow for sp9C1 in 300mM KCl at pH 8. **E** Two perpendicular side views of the pH 8/KCl map, before (top) and after (bottom) map sharpening. Maps are colored according to local resolution (FSC = 0.5) as indicated in the color key. **F** Summary of single particle reconstruction of sp9C1 at pH 8 in presence of 300mM NaCl

**Extended Figure 2.**
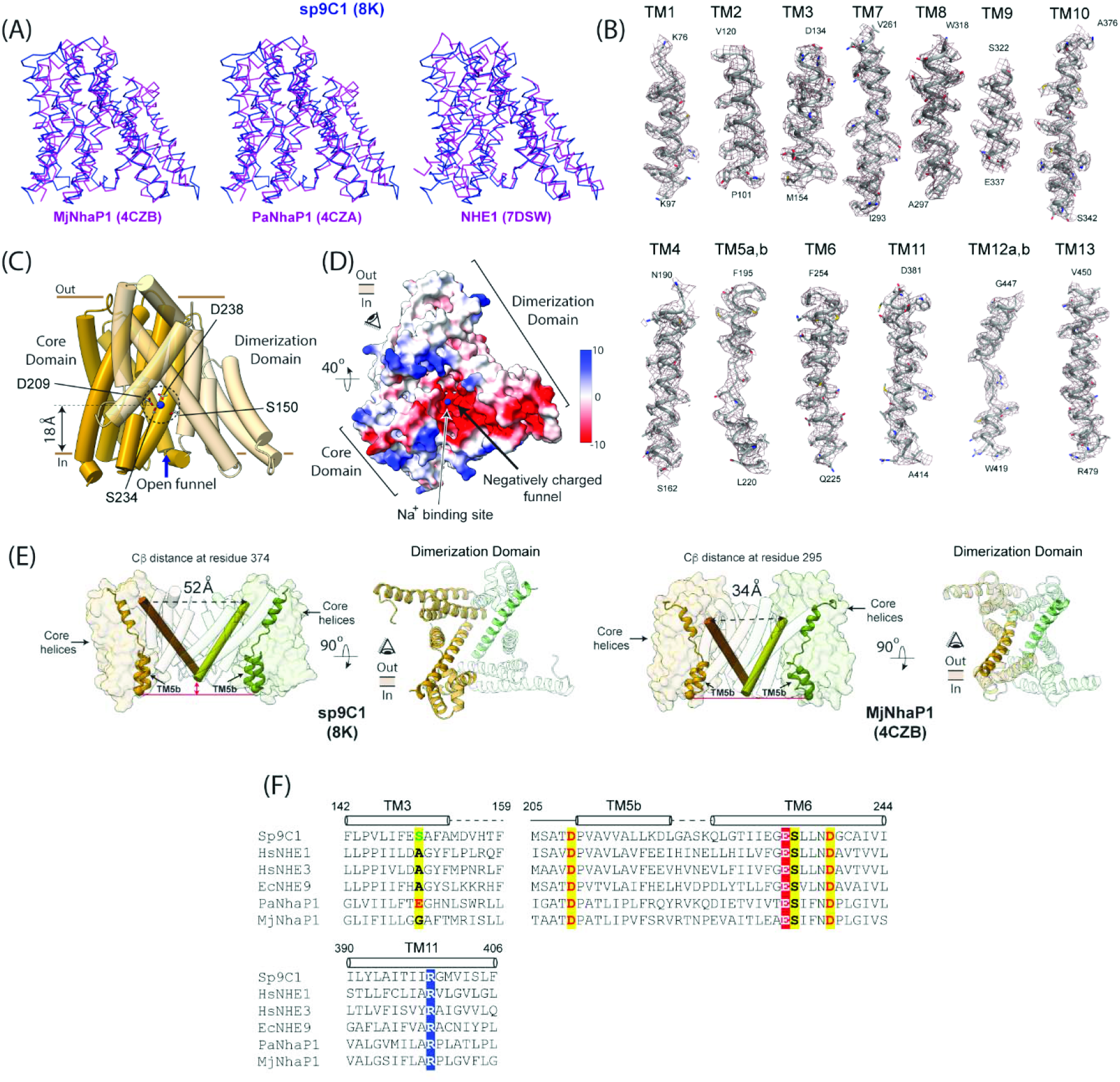
The Ion-Exchange Domain of sp9C1. **A** Structural comparison of a single protomer of the ion exchange domain of sp9C1 (pH 8/KCl), in blue, with those of three representative SLC9s obtained using X-ray crystallography (MjNhaP1 and PaNhaP1) and single-particle cryoEM (NHE1 in the inward-open state), in magenta. **B** Densities of the 13 transmembrane helices of the ED as observed in the pH 8/KCl sharpened map comparing the model to the map. **C** Side view of the ED showing the helices of the dimerization domain in transparent yellow and core domain in dark yellow. An open funnel connects the cytoplasm to the ion binding site deep in the transmembrane domain. Residues at the principal ion binding site are shown in sticks. The blue sphere is the mass center of the side chains of the 4 annotated residues and coincides with the putative substrate binding locus. **D** Bottom-up view of the ED showing the negative electrostatic surface of the cytoplasmic funnel with the putative sodium binding site at its tip. **E** Comparison of the ED dimer of sp9C1 (side and top-down views on the left) and MjNhaP1 (side and top-down views on the left). TM10s of the two subunits which line the inter-subunit cavity are shown as brown and green cylinders in side-view representations. The exterior ends of TM10 are significantly further apart in sp9C1 than in MjNhaP1 owing to the wider inter-subunit cavity. TM5 (and the intracellular end of TM6) of the 2 subunits are shown in yellow and green cartoon representations. In sp9C1, the intracellular tip of TM5b is ∼6Å further below the intracellular ends of the dimerization domain (at the level of TM10) as compared to that in MjNhaP1. The inner surface of sp9C1 ED-dimer is thus significantly more concave. **F** Sequence alignment of key regions of the ED of different SLC9s, highlighting the ion binding sites in yellow. Two other highly conserved residues, R399 and E233, are also highlighted in blue and red respectively.

**Extended Figure 3.**
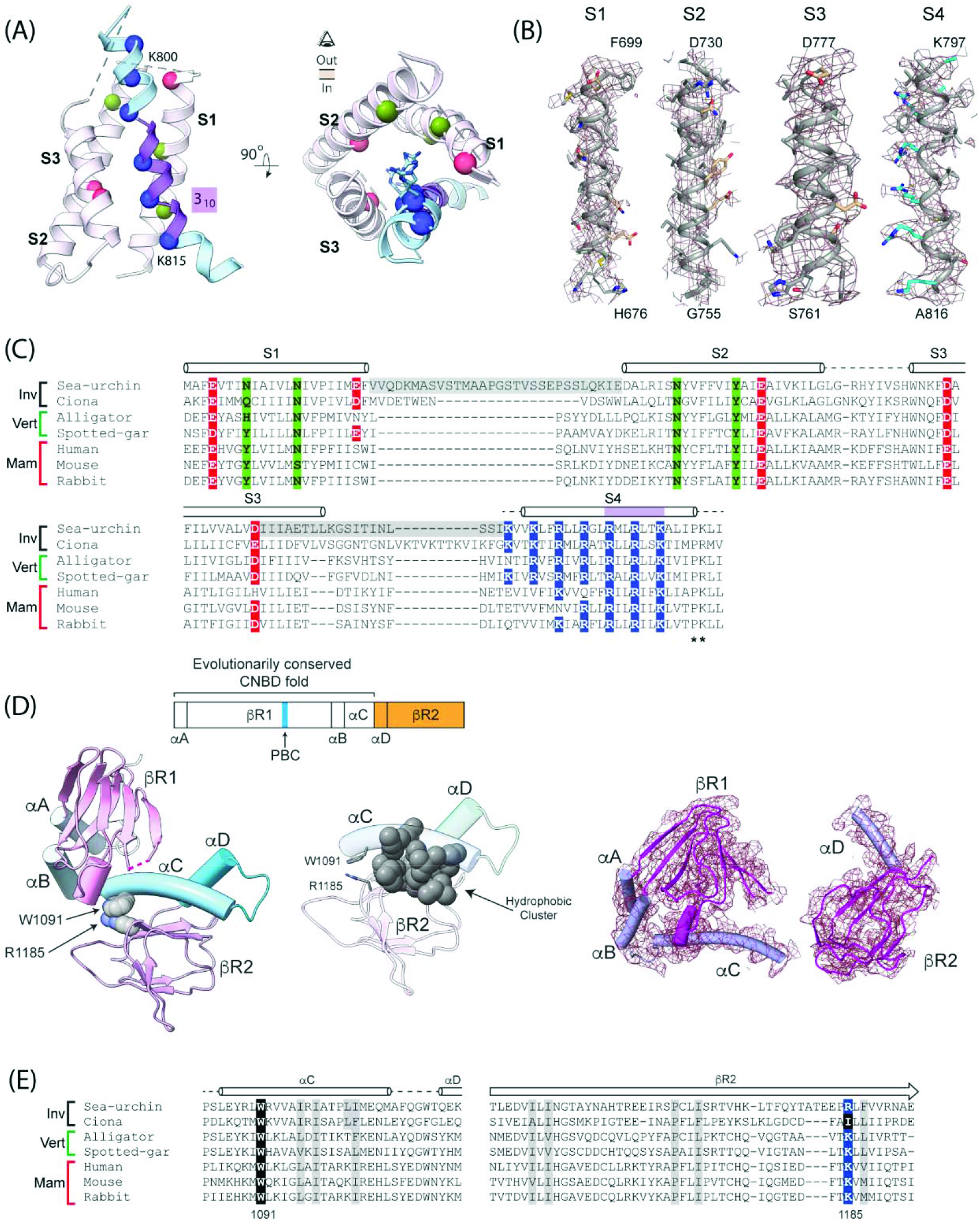
Regulatory domains of sp9C1. **A** Side-and top-down views of voltage-sensing domain. Cα atoms pf positively charged residues on S4 are shown in blue spheres. The 3_10_ helical segment of S4 is colored in purple while the rest is alpha helical shown in light blue. The positions of polar/negatively charged residues on S1-S3, which putatively stabilize the S4 charges, are shown in green and red spheres. **B** Density maps of the four helices of the VSD. Side chains of the S4 charges are colored in blue/cyan. **C** Sequence alignment of the voltage-sensing domain of invertebrate (Inv), vertebrate (Vert) and mammalian (Mam) SLC9C1s. Residues are highlighted as in (A). The extracellular ends of S1 and S2 and S3 and S4, which are missing in density, are highlighted in grey. **D** Topology, structure and density map of the cytoplasmic domain of sp9C1 showing the CNBD and βR2 domains. PBC (highlighted in blue in the topology) refers to the ‘phosphate binding cassette’, an evolutionarily conserved motif featuring a cluster of positively charged residues that binds the phosphate group of cyclic nucleotides. A conserved pair of residues (W1091 and R1185) and hydrophobic residues at the interface between αC and βR2 are shown in spheres. Red dotted line indicates a short loop in the βR1 domain that is missing in density. **E** Sequence alignment of αC to βR2 regions of SLC9C1 in various SLC9s showing the interfacial residues (depicted as spheres in (D)), with the W1091 and R1185 sites highlighted in black and blue respectively.

**Extended Figure 4.**
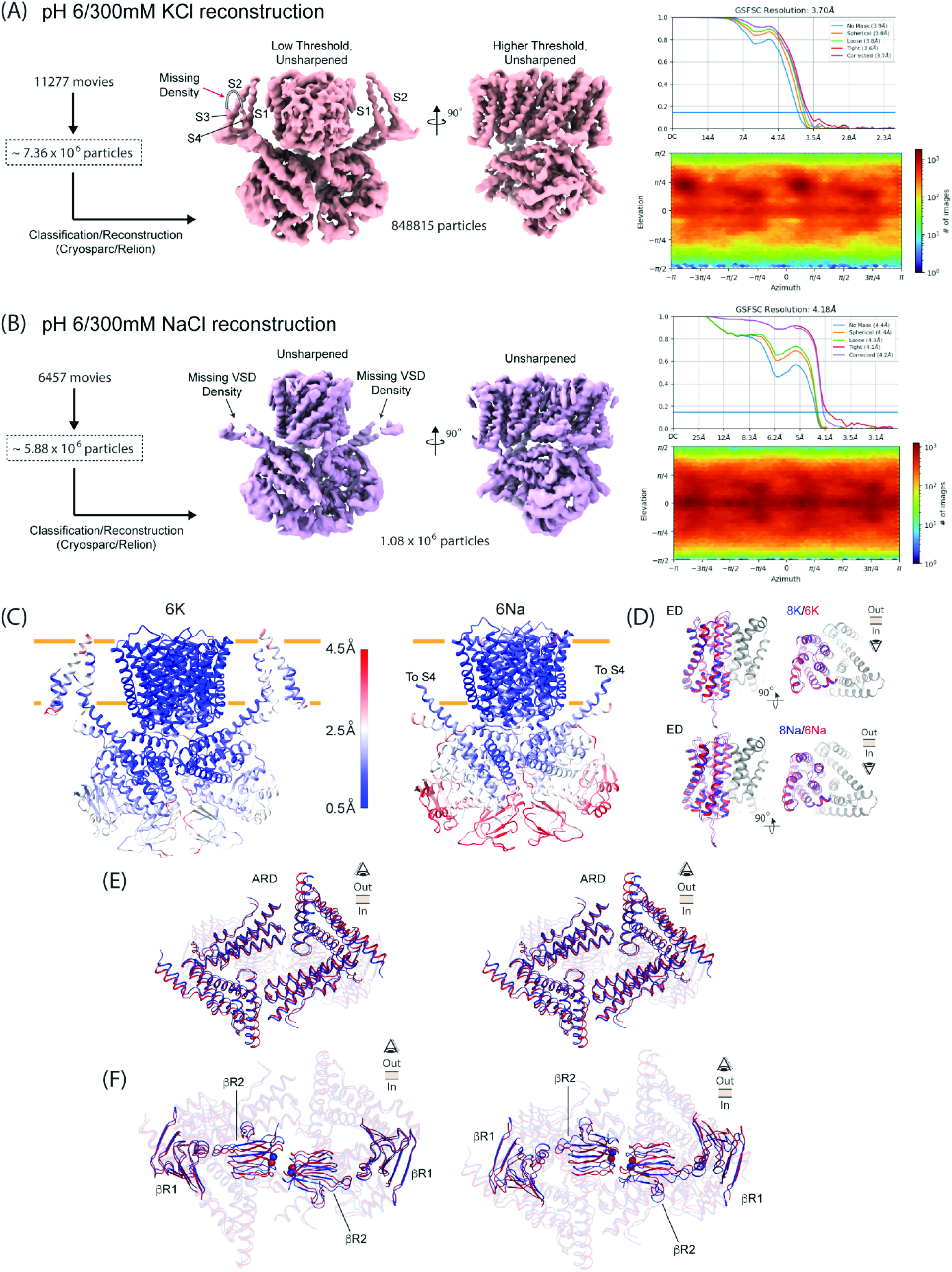
Single particle reconstruction of WT sp9C1 in pH 6. **A, B** Summary of single particle reconstruction of sp9C1 at pH 6, obtained in 300mM KCl (A) or 300mM NaCl (B). The density for the extracellular ends of S3 and S4 helices of the VSD are virtually absent in (A) while in (B), the density for the entire VSD averages out on account of structural heterogeneity. **C** Structural models of sp9C1 at pH 6, obtained in 300mM KCl (left) or 300mM NaCl, colored according to Cα displacements relative to pH 8/KCl or pH 8/NaCl structures, respectively. **D, E, F** Structural comparison of the ED (D), ARD (E) and the two β-rolls (F), between pH 8 (blue) and pH 6 (red), in 300mM KCl or NaCl, as indicated. Small pH dependent displacements are observed at the level of βR2 in presence of KCl or NaCl (F). Blue and red spheres indicate the Cα of residue H1172 in pH 8 and pH 6 structures respectively.

**Extended Figure 5.**
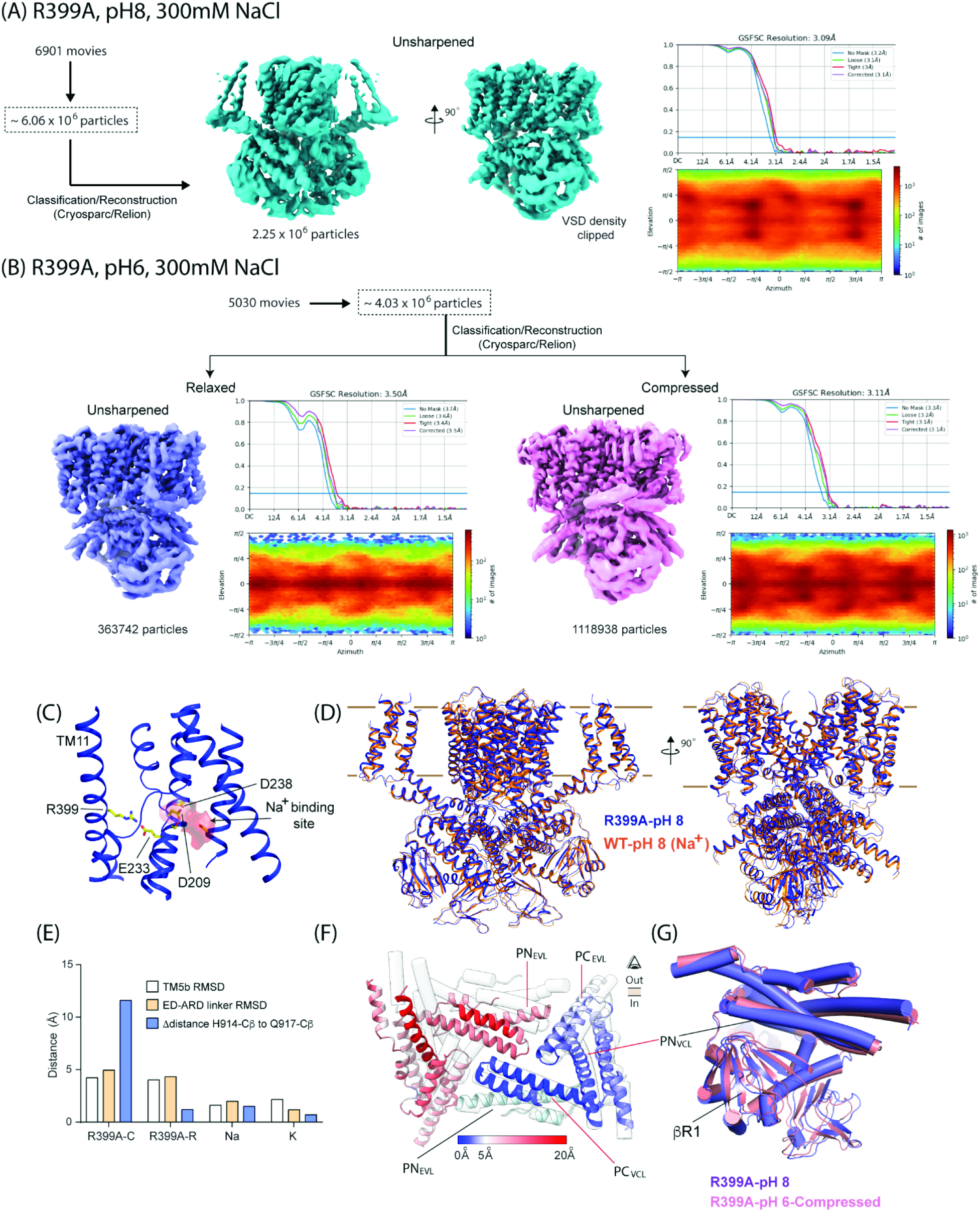
Single particle reconstruction of R399A mutant of sp9C1. **A, B** Summary of single particle reconstruction of R399A-sp9C1 at pH 8 (A) and pH 6 (B). At pH 6, the number of particles contributing to the Compressed class is ∼3-fold higher than those contributing to the Relaxed class. Much like WT, the density for the VSD in both pH 6 classes is virtually absent. **C** In WT sp9C1, R399 forms a salt-bridge interaction with E233. The salt bridge is close to the ion binding site in the inward-open state of the ED. **D** Structural comparison of R399A-sp9C1 (blue) and WT-sp9C1 (orange) at pH 8. The two structures superposed well with an overall RMSD of 1.45Å. **E** Changes in key structural parameters between pH 8 and pH 6 for R399A-Compressed class (R399A-C, group 1), R399A-Relaxed class (R399A-R group 2), WT-sp9C1 in presence of NaCl (Na, group 3) and KCl (K, group 4). The pH 8 structures, used for RMSD and distance calculations, were R399A-pH 8 (for groups 1 and 2), pH8/NaCl (group 3) and pH 8/KCl (group 4). **F** Structural superposition of ARDs in R399A-pH 6 Compressed Class (colored cartoon) and R399A-pH 8 (transparent tubes). The two ARDs are aligned at using PN_EVL_ of single subunit (marked with black line). Cartoon colors represents the Cα displacements between the two structures. One side of the ARD shows limited displacements with respect to the other. *Left*, ARD segments belonging to the same subunit are marked with the red line. **G** Side view ARD-βR1 interface of R399A-pH 6 Compressed Class (pink) and R399A-pH 8 (purple) aligned at the level of the ARD showing that the ARD (PN_VCL_) and βR1 interface remains preserved.

**Extended Figure 6.**
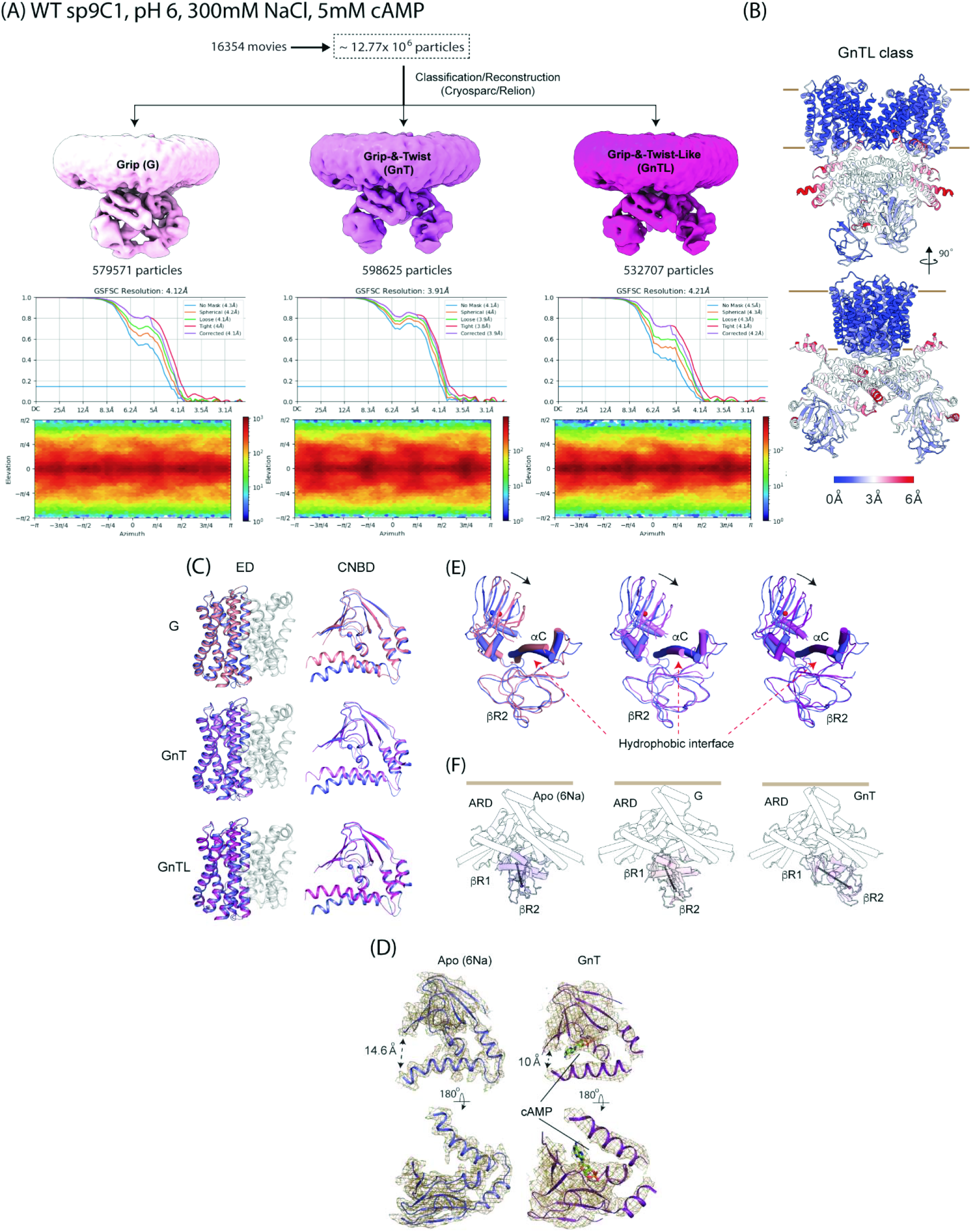
Single particle reconstruction of cAMP bound sp9C1. **A** Summary of single particle reconstruction of WT sp9C1 in pH 6, with 300mM NaCl and 5mM cAMP showing the 3 structural classes obtained and their final resolutions. Maps are shown at a low contour level to highlight the difference in the cytosolic domains. **B** Model of the GnTL class in two perpendicular side views, colored according to Cα displacements relative to the GnT class, showing the differences in the ARD and C-terminal domains between the two structures. **C** Comparison of the ED and CNBD between the Apo (pH 6/NaCl) structure (in blue) and the 3 cAMP conformations, G (light pink), GnT (pink) and GnTL (dark pink). The spheres in the CNBD indicate the Cα atom of R1053 that is involved in cAMP binding. The dimerization domain helices are shown in gray. **D** Density maps of the CNBD in the Apo (pH 6/NaCl) and cAMP bound (GnT) states, showing the cAMP density. **E** Structural superposition of the cytoplasmic domains (CNBD and βR2) between Apo and cAMP bounds forms, colored as in (C). Blue and red spheres indicate the center of masses of βR1 in the Apo and cAMP bound states and arrow shows the movement of βR1. **F** Side view of the full ARD (white tubes) and the cytoplasmic domain of a single subunit in Apo, G and GnT states. In each case the solid black line connects the center of masses of βR1 and βR2. The solid brown lines indicate the inner leaflet of the membrane.

## Notes

### Competing Interest Statement

The authors have declared no competing interest.

