## Supplementary Information for "Architecture and rearrangements of a sperm-specific Na^+^/H^+^ exchanger"

**Supplementary Table I. Cryo-EM data collection, refinement and validation statistics for WT sp9C1 reconstructions**

|  | **pH 8 Na** | **pH 8 K** | **pH 6 Na** | **pH 6 K** |
| --- | --- | --- | --- | --- |
| EMDB # | 41965 | 41964 | 41991 | 41990 |
| PDB-ID | 8U71 | 8U6Z | 8U7S | 8U7R |
| **Data collection and processing** |  |  |  |  |
| Magnification | 81000 | 64000 | 64000 | 81000 |
| Voltage (kV) | 300 | 300 | 300 | 300 |
| Electron exposure (e–/Å^2^) | 66 | 65 | 65 | 66 |
| Defocus range (μm) | -0.9 to -1.7 | -0.9 to -1.7 | -0.9 to -1.7 | -0.9 to -1.7 |
| Pixel size (Å) (super-resolution)  (Binned by 2 during data processing) | 0.55 | 0.69 | 0.69 | 0.55 |
| Symmetry imposed | C2 | C2 | C2 | C2 |
| Initial particle images (no. x 10^6^) | 2.58 | 3.89 | 5.88 | 7.36 |
| Final particle images (no.) | 150270 | 563022 | 1088719 | 848815 |
| Map resolution (Å) | 3.9 | 3.8 | 4.2 | 3.7 |
| FSC threshold | 0.143 | 0.143 | 0.143 | 0.143 |
| **Refinement** |  |  |  |  |
| Model resolution (Å) | 4.1 | 4.0 | 4.2 | 3.9 |
| FSC threshold | 0.5 | 0.5 | 0.5 | 0.5 |
| Map sharpening *B* factor (Å^2^) | -126 | -186 | -175 | -168 |
| Model Composition |  |  |  |  |
| Non-Hydrogen atoms | 16033 | 16046 | 14630 | 15601 |
| Protein Residues | 1990 | 1998 | 1816 | 1971 |
| Ligand | 7 | 7 | 7 | 7 |
| ***B-factors (Å^2^)*** |  |  |  |  |
| Protein | 79.97 | 55.02 | 53.33 | 41.56 |
| Ligand | 58.55 | 44.98 | 44.13 | 35.09 |
| ***R.m.s Deviations*** |  |  |  |  |
| Bond lengths (Å) | 0.004 | 0.004 | 0.003 | 0.003 |
| Bond Angles (°) | 0.762 | 0.672 | 0.711 | 0.643 |
| ***Validation*** |  |  |  |  |
| MolProbity Score | 1.55 | 1.42 | 1.39 | 1.35 |
| Clash score | 7.21 | 4.97 | 5.89 | 6.34 |
| Poor rotamer (%) | 0.18 | 0.00 | 0.06 | 0.13 |
| ***RamaChandran*** |  |  |  |  |
| Favored (%) | 97.15 | 97.11 | 97.67 | 98.15 |
| Allowed (%) | 2.85 | 2.89 | 2.33 | 1.85 |
| Outlier (%) | 0.00 | 0.00 | 0.00 | 0.00 |

**Supplementary Table II. Cryo-EM data collection, refinement and validation statistics for R399A-sp9C1 reconstructions**

|  | **pH 8 R399A** | **pH 6 R399A Relaxed** | **pH 6 R399A Compressed** |
| --- | --- | --- | --- |
| EMDB # | 41967 | 41979 | 41969 |
| PDB-ID | 8U73 | 8U7D | 8U75 |
| **Data collection and processing** |  |  |  |
| Magnification | 64000 | 64000 | 64000 |
| Voltage (kV) | 300 | 300 | 300 |
| Electron exposure (e–/Å^2^) | 66 | 66 | 66 |
| Defocus range (μm) | -0.9 to -1.7 | -0.9 to -1.7 | -0.9 to -1.7 |
| Pixel size (Å) (super-resolution) | 0.68 | 0.68 | 0.68 |
| Symmetry imposed | C2 | C2 | C2 |
| Initial particle images (no. x 10^6^) | 6.06 | 4.03 | 4.03 |
| Final particle images (no.) | 2251538 | 363742 | 1118938 |
| Map resolution (Å) | 3.1 | 3.5 | 3.1 |
| FSC threshold | 0.143 | 0.143 | 0.143 |
| **Refinement** |  |  |  |
| Model resolution (Å) | 3.2 | 3.6 | 3.2 |
| FSC threshold | 0.5 | 0.5 | 0.5 |
| Map sharpening *B* factor (Å^2^) | -184 | -144.6 | -154.6 |
| Model Composition |  |  |  |
| Non-Hydrogen atoms | 16199 | 14924 | 14656 |
| Protein Residues | 2006 | 1846 | 1814 |
| Ligand | 7 | 7 | 7 |
| ***B-factors (Å^2^)*** |  |  |  |
| Protein | 16.68 | 70.04 | 38.57 |
| Ligand | 3.45 | 53.32 | 20.88 |
| ***R.m.s Deviations*** |  |  |  |
| Bond lengths (Å) | 0.003 | 0.004 | 0.004 |
| Bond Angles (°) | 0.690 | 0.592 | 0.642 |
| ***Validation*** |  |  |  |
| MolProbity Score | 1.40 | 1.42 | 1.35 |
| Clash score | 7.19 | 5.54 | 5.7 |
| Poor rotamer (%) | 0.12 | 0.13 | 0.06 |
| ***RamaChandran*** |  |  |  |
| Favored (%) | 98.08 | 97.37 | 97.83 |
| Allowed (%) | 1.92 | 2.63 | 2.17 |
| Outlier (%) | 0.00 | 0.00 | 0.00 |

**Supplementary Table III. Cryo-EM data collection, refinement and validation statistics for WT sp9C1 in cAMP bound states**

|  | **G** | **GnT** | **GnTL** |
| --- | --- | --- | --- |
| EMDB # | 41984 | 41987 | 41988 |
| PDB-ID | 8U7K | 8U7O | 8U7P |
| **Data collection and processing** |  |  |  |
| Magnification | 64000 | 64000 | 64000 |
| Voltage (kV) | 300 | 300 | 300 |
| Electron exposure (e–/Å^2^) | 66 | 66 | 66 |
| Defocus range (μm) | -0.9 to -1.7 | -0.9 to -1.7 | -0.9 to -1.7 |
| Pixel size (Å)  (Binned by 2 during data processing) | 0.69 | 0.69 | 0.69 |
| Symmetry imposed | C2 | C2 | C2 |
| Initial particle images (no. x 10^6^) | 12.77 | 12.77 | 12.77 |
| Final particle images (no.) | 579571 | 598625 | 532707 |
| Map resolution (Å) | 4.1 | 3.9 | 4.2 |
| FSC threshold | 0.143 | 0.143 | 0.143 |
| **Refinement** |  |  |  |
| Model resolution (Å) | 4.2 | 4.0 | 4.2 |
| FSC threshold | 0.5 | 0.5 | 0.5 |
| Map sharpening *B* factor (Å^2^) | -249 | -214.6 | -258.4 |
| Model Composition |  |  |  |
| Non-Hydrogen atoms | 14147 | 14929 | 14086 |
| Protein Residues | 1812 | 1959 | 1808 |
| Ligand | 9 | 9 | 9 |
| ***B-factors (Å^2^)*** |  |  |  |
| Protein | 124.11 | 85.89 | 172.37 |
| Ligand | 112.30 | 68.91 | 119.54 |
| ***R.m.s Deviations*** |  |  |  |
| Bond lengths (Å) | 0.005 | 0.002 | 0.005 |
| Bond Angles (°) | 0.888 | 0.606 | 1.227 |
| ***Validation*** |  |  |  |
| MolProbity Score | 1.45 | 1.29 | 1.47 |
| Clash score | 8.29 | 5.42 | 8.67 |
| Poor rotamer (%) | 0.00 | 0.07 | 0.00 |
| ***RamaChandran*** |  |  |  |
| Favored (%) | 98.27 | 98.29 | 98.44 |
| Allowed (%) | 1.73 | 1.71 | 1.56 |
| Outlier (%) | 0.00 | 0.00 | 0.00 |

**SUPPLEMENTARY LEGENDS**

**Supplementary Figure 1. Model to map validation statistics for sp9C1 reconstructions.** Model to map FSCs of the 10 reconstructions of sp9C1 (and R399A mutant) obtained from the real space refinement module in Phenix. Results show the refinement against the final sharpened map.

**Supplementary Video 1. Architecture of sp9C1**. Different views of sp9C1 (pH 8/K model) showing the arrangement of its different structural elements at a single subunit level and how they are organized in the context of sp9C1 dimers. Outline marks the surface of the model. The structural elements demarcated as “Paddle” indicate the N and C terminal paddles of the EVL and VCL which together form the ARD. VSD, CNBD and B-R2 implies the voltage-sending domain, cyclic nucleotide binding domain and the β-R2 domains respectively.

**Supplementary Video 2. pH dependent transitions of the R399A mutant of sp9C1.** Morph showing the transitions from pH 8 to the pH 6 compressed class via the intermediate pH 6 relaxed class, in different views, as observed in the R399A mutant (in presence of NaCl). Intracellular ends of TM5b, TM13 and the short TM13-ARD linker (marked in red) move between pH 8 and pH 6 relaxed state but during this transition there are limited changes at the level of the ARD. The ARD rearranges between the pH 6 relaxed and compressed states. VSDs are not modeled in the pH 6 states and are not featured in the morphs.

**Supplementary Video 3. cAMP dependent transitions of sp9C1.** Morph showing the cAMP dependent conformational dynamics of sp9C1. Initial morphs depict the transitions from the pH 6/Na model to the Grip (G) state and finally to the Grip-and-Twist (GnT) state. Snapshots of the CNBD (βR1) and ARD (PN_VCL_) in the various states highlight the reorganization of the intra-subunit interface accompanying cAMP binding. The ARD reorganizes in the G and GnT. These morphs do not feature the VSD as they are not modeled in the 6/Na and G states. The final morph between pH 6/K and GnT states, where the VSD helices are defined, show the lateral displacement of the VSDs, accompanying ARD rearrangement in response to cAMP binding.
